# Social defeat induces REM sleep fragmentation through the PFC-VLPO pathway

**DOI:** 10.1101/2022.12.27.522036

**Authors:** M. Chouvaeff, S. Bagur, L. Mace, T. Gallopin, K. Benchenane

## Abstract

The brainstem and hypothalamic structures implementing the daily cycles of wake, REM and NREM sleep have now been identified in remarkable detail. However, sleep structure dynamically adapts to environmental stressors, likely requiring top-down cortical feedback that is as yet unidentified. Here, we investigate the role of projections from prefrontal cortex (PFC), a key hub in stress regulation, to the ventro-lateral preoptic areas (VLPO), a master regulator of sleep states. Using *ex vivo* optogenetics, we demonstrate that activation of PFC terminals induces monosynaptic excitatory glutamatergic currents in VLPO NREM-promoting neurons. *In vivo*, activation of PFC-VLPO projections interrupts ongoing REM in favour of NREM, leading to fragmented REM bouts. Remarkably, chemogenetic inhibition of PFC-VLPO projections has no effect in baseline conditions but it blocks the REM fragmentation induced by Social Defeat Stress. Therefore, the PFC-VLPO pathway provides a top-down regulation specifically recruited in stressful conditions to induce short, fragmented REM bouts and favor NREM sleep.

## INTRODUCTION

Mammals alternate between two sleep phases, NREM and REM sleep. In NREM sleep, brain activity is dominated by high-voltage, low frequency activity, while in REM sleep, the electroencephalogram (EEG) is more similar to wakefulness, with higher theta power, but a complete lack of muscle tone (Dement, 1958; Jouvet, 1962). Many studies have identified the neural circuits involved in the baseline initiation and maintenance of these different sleep states (Scammell et al., 2017). Moreover, sleep state can be influenced by a variety of regulatory pathways, including circadian, homeostatic, temperature, and metabolic regulation (Saper et al., 2010; Scammell et al., 2017; Luppi and Fort, 2019). This dynamic, context-dependent regulation is still poorly understood.

The level of stress experienced by humans or animals is one of the strongest disruptors of sleep. Stress-induced sleep perturbations are characteristics of many psychiatric disorders such as depression, generalized anxiety and post-traumatic stress disorder (PTSD) (Papadimitriou and Linkowski, 2005). Although the effects of stress on sleep can lead to serious health problems, we lack a clear picture of how stress impacts sleep. Indeed, depending on the type of stress, its duration of exposure, and the circadian period at which it occurs, the effects of stress on sleep can vary significantly, including fragmentation caused by multiple awakenings, a reduction or an increase of NREM or REM sleep (Pawlyk et al., 2008; Sanford et al., 2015; Antila et al., 2022; Yu et al., 2022). This complexity highlights the need to identify the different neural circuits that regulate the sleep response to stress in order to parse their contributions to specific aspects of sleep architecture.

Whereas the baseline regulators of sleep states are found in the brainstem and hypothalamus, it is likely that the flexible and contextual adaptations linked with stress will rely on top-down input from higher-level structures. The most likely candidate is the infralimbic (IL) region of the prefrontal cortex (PFC) which is ideally positioned to mediate between stressors and sleep. On the one hand, it is heavily interconnected with limbic structures and plays a significant role in modulating fear-related behaviors, especially those related to emotional response regulation (Sullivan and Gratton, 2002). On the other hand, the IL projects to the ventrolateral preoptic area (VLPO) (Chou et al., 2002) which directly controls NREM sleep initiation and maintenance (Sherin et al., 1996, 1998; Szymusiak et al., 1998; Lu et al., 2000; Takahashi et al., 2009; Chung et al., 2017; Kroeger et al., 2018). Moreover, ventromedial PFC lesions, including the IL, increase REM sleep and sleep fragmentation (Chang et al., 2014). Interestingly, stress induced by cage-exchange leads to an activation of neurons in the infralimbic (IL) region of the PFC as well as NREM sleep promoting neurons in the ventrolateral preoptic nucleus (VLPO) (Cano et al., 2008).

These results suggest that the IL projections to the VLPO could mediate the link between stress and sleep architecture. To date, how cortical projections modify the activity of this sleep-promoting structure remains unclear and no direct causal link has been established. We therefore tested the hypothesis that the PFC directly influences sleep-promoting neurons and participates in the regulation of sleep state transitions under stressful conditions. Using circuit tracing and optogenetic approaches in mice, both *in vivo* and *ex vivo*, we identified the synaptic mechanisms by which VLPO activity is modulated by the PFC, leading to a fragmentation of REM sleep. Using chemogenetic inhibition, we additionally investigated whether activation of this cortical pathway contributes to the reduction of REM sleep in mice when they sleep in dangerous environments.

## MATERIAL AND METHODS

### Animals and housing

Adult male C57Bl/6J (6–15 week-old) and outbred male Swiss CD1 mice (9–16 week-old) were used in this study (Centre d’élevage R. Janvier, Le Genest St. Isle, France). All behavioural experiments were conducted in accordance with the official European guidelines for the care and use of laboratory animals (86/609/EEC) and in accordance with the Policies of the French Ethics Committee. Experiments at the ESPCI were conducted in an animal facility that has been fully accredited by the French Direction of Veterinary Services (D75-05-14). An ethical committee and the Ministry of Research approved the animal experiments carried out in this study (APAFIS#240242020021014469086). We made every effort to limit the number of animals and their suffering as much as possible. The mice were housed at 22 ± 2 °C and on a 12 h light/dark cycle (08:00 - 20:00 light), with food and water available ad libitum.

### Viral vectors

For the expression of the photo-sensitive cation channel Channelrhodopsin-2 (ChR2) in PFC neurons, an adeno-associated virus (AAV9-pCaMKII-Chr2-mCherry, UNC Vector Core, USA) was injected in the Infralimbic (IL) / Prelimbic (PrL) cortex. Control mice were injected with a virus lacking the ChannelRhodopsin sequences (AAV9-pCaMKII-mCherry, UNC Vector Core, USA). For Cre-recombinase-dependent expression of the inhibitory designer receptor hM4D(Gi) in the PFC neurons projecting to the VLPO, we used the adeno-associated viral vector AAV-hSyn-DIO-hM4D(Gi)-mCherry (Viral Vector Facility, Zurich, Switzerland). The Cre-recombinase was retrogradely brought via an injection of the retro-adeno-associated viral vector rAAV-hSyn-EGFP-Cre (Viral Vector Facility, Zurich, Switzerland) in the VLPO. This strategy allowed us to modulate specifically the activity of the PFC neurons projecting to the VLPO.

### Stereotaxic viral injections

6-week-old mice were anesthetized in an induction chamber with 3 % isoflurane. Following anesthetic induction mice received analgesia (buprenorphine, 0,1mg/kg, subcutaneous (s.c.) injection) and were placed in a stereotaxic frame (David Kopf Instruments, USA). Throughout the whole procedure, the isoflurane concentration was maintained at 1.5 - 2%. Core body temperature was maintained at 37°C with a heat pad. A local subcutaneous injection of lidocaine (4-5mg/kg) was performed before the skin incision. Bilateral craniotomies were performed using a dremel above structures of interest, IL/PrL (AP: +1.75, ML: ± 0.5, DV: - 2.1) and VLPO (AP: + 0.7, ML: ± 0.7, DV: - 5.1). A Hamilton syringe was lowered and maintained for 5 min before the beginning of viral injection (150 nL, 15 nL/min) and for an additional 5 min after injection to avoid vector reflux. The injection was performed using a programmable pump (LEGATO 130, KDScientific). The incision was sutured and disinfected. Mice received subcutaneous injection of Metacam (1m g/kg) and/or Buprenorphine (0.1 mg/kg) and recovered from surgery in individual cages on a heated mat for 24 h. They were then maintained at an ambient temperature of 22°C with food and water available ad libitum.

### Slice preparation for electrophysiological recordings

For recordings performed in the VLPO or in the PFC, mice aged 7 to 8 weeks were deeply anesthetized with ketamine (1%) and xylazine (1‰) and were perfused intracardially with an ice-cold sucrose solution containing (in mM): Sucrose (230), CaCl2 (0,5), glucose (10), KCl (2.5), MgSO4 (10), NaHCO3 (26); NaH2PO4 (1.25), kynurenate (3). The animals were decapitated and brains were quickly removed in the same ice-cold solution. Coronal slices (250 µm thick) were cut with a vibrating microtome (VT2000S; Leica), collected and maintained for 10 minutes in a chamber containing artificial Cerebro-Spinal Fluid (aCSF) warmed to 36°C containing (in mM): 130 NaCl; 5 KCl; 2.4 CaCl2; 20 NaHCO3; 1.25 KH2PO4; 1.3 MgSO4; 10 D-glucose; and 15 sucrose ; kynurenate (1) (pH = 7.35). Slices were then transferred in another chamber containing the same previous aCSF solution without kynurenate and at room temperature. A recovery period of 60 minutes was then observed before slices were transferred to the recording chamber, where they were maintained immersed and continuously supervised at 1-2 ml/min with the same aCSF saturated with 95% oxygen and 5% carbon dioxide.

### Whole cell Patch-Clamp recordings

Individual slices were submerged in a recording chamber, placed on the stage of an Axioskop 2FS microscope (Carl Zeiss), equipped with Dodt gradient contrast optics (Luigs & Neuman), and an infrared CCD camera (CoolSNAP HQ2; Roper Scientific, Germany). Neurons in the slices were visualized by using infra-red (IR) videomicroscopy. Pipettes (4 to 6 MΩ) were pulled from borosillicate capillaries (1,5 mm o.d., 0,86 mm i.d., Harvard Appartus, France) and filled with 8 µl of autoclaved internal solution containing 144 mM K-gluconate, 1 mM MgCl2, 0.2 mM EGTA, 10 mM HEPES, pH 7.2 (285/295 mOsm) and were attached to a micromanipulator (Luigs er Neuman). Because the intracellular solution used was depleted in chloride, the equilibrium potential of this ion was: ECl ≈ -90mV. Thus, potential GABAergic currents appear outward at potentials more depolarized than ECl whereas cationic currents are inward. All electrophysiological experiments were performed with a MultiClamp700B (Axon Instruments) amplifier connected to an acquisition board (Digidata 1440; Axon Instruments) attached to a computer running pCLAMP software (Axon Instruments). Axonal fibers from the PFC expressing ChR2 were stimulated in the VLPO by pulses of 470 nm light of 5ms every 5s for 1 to 5 min using a diode laser (pE-2, Coolled). The currents evoked by the photo-stimulations were recorded in voltage-clamp configuration and characterized using the application of the following antagonists: kynurenate (1mM; Sigma); tetrodotoxin (TTX, 1µM; Sigma); 4 aminopyridine (4-AP, 100µM; Sigma). The currents recorded at the same imposed potential were averaged for each cell to perform the quantification. Norepinephrine (NA, 100µM, Sigma) was also bath applied to identify the potential induced inhibition characteristic of VLPO sleep promoting neurons (Gallopin et al., 2000, 2005; Matsuo et al., 2003; Saint-Mleux, 2004; Liu et al., 2010).

### Stereotaxic electrode and optical fibers implantations

Two weeks following vector injections, mice were anesthetized according to the same procedure as presented above. Isoflurane anesthesia was maintained throughout the surgery between 1.5 to 2% by inhalation and placed on a stereotaxic frame. Electrodes (tungsten wires) were bilaterally implanted in the Olfactory Bulb (OB) (AP: + 4.3, ML: ± 0.5, DV: -1.5), in the CA1 hippocampal layer (AP: - 2.2, ML: ± 2.0, DV: - 1.0) and the Prefrontal cortex (PFC) (AP: + 1.95, ML: ± 0.5, DV: -1.5). A reference electrode was placed on the surface of the cerebellum. A single or two wires hooked into the nuchal muscles were inserted in two out of six optogenetic mice, as well as in all chemogenetic mice. During recovery from surgery and during all experiments, mice received food and water ad libitum. Recordings began 1 week after surgery. In addition to the electrodes implantation, the optogenetic mice groups were also bilaterally implanted with optical fibers (0.22 mm diameter, spaced 1.4 mm, Doric Lenses Inc, Quebec, Canada) during the same surgery. Fibers were lowered just above the VLPO (AP: + 0.65, ML: ± 0.7, DV: −5.1). On each optical fiber, tetrodes formed by twisting Nichrome wires (0.001” bare) were glued in order to record the LFP signals from the VLPO. At the end of implantation, the electrodes and optical fibers were fixed to the skull with wax and dental cement (SuperBond C & B). The bilateral optical fibers and the electrodes were then connected to an EIB16 card held on the surface of the skull by parafilm and dental cement. Brain activity was recorded after at least one week of recovery.

### Optogenetic protocol

All experiments used 473nm light, delivered using a laser (Laserglow Technology) connected to a rotating connector (Doric Lense). Patch cords are then connected to optical fibers (numerical aperture 0.37) on the animal’s head. Mice were habituated to the recording cable for at least 2 days before starting the recordings. Recordings were made from around 8:30 a.m. to 6:30 p.m. hoto-stimulations (20 Hz, 30s, 4 mW at the 200 μm fiber output) were only delivered during sleep by automatically triggering them when the animal’s activity measured with a camera (FLIR SC325) fell below a threshold for at least 1 minute. The minimal inter-stimulations-interval was set at 1 minute.

### Chemogenetic protocol

The clozapine-N-oxide (CNO, agonist at hM4D(Gi) receptors) was purchased from HelloBio and stored in a freezer at -30°C until use. Before each experiment CNO was diluted to final concentrations and mice were injected intraperitoneally with either CNO (2.5mg/kg) or a vehicle control solution (Saline, NaCl 9g/l) at 1pm. Recordings were made from around 8:30 a.m. to 6:30 p.m.

### Social defeat stress (SDS) protocole

The social defeat stress procedure used in this study was adapted from a previous published method (Henderson et al., 2017). The experiment started at 9am. First, the intruder (experimental mouse) was introduced into the home cage of an unfamiliar, aggressive CD1 resident mouse during 5 min for physical interactions. If the conflicts were too intense, the mice were briefly separated for a few seconds to avoid wounds and damages to the implant. In a second step, the two mice were separated by a transparent and perforated partition placed in the middle of the CD1 mouse home cage for 20 minutes. During this step, the experimental mouse remained in sensory contact (olfactory, visual and auditory) with the CD1 mouse. In a final step, a second sensory exposure of 20 minutes was repeated in the experimental mouse’s home cage to reinforce the association between its home cage and the stressful experience. During the stress procedure the experimental mouse usually shows clear submissive posture and freezing behavior. The CD1 mouse was then removed from the cage and the experimental mouse was treated either with saline or CNO (2.5mg/kg, i.p). The mouse was then connected to the recording cable and allowed to sleep for the rest of the day (in its home cage with the partition). The sleep sessions usually started around 10 am up to 6:30 pm.

### *In vivo* Electrophysiological recordings

Signals from all electrodes were recorded using an Intan Technologies amplifier chip (RHD2216, sampling rate 20 KHz). Local Fields Potentials (LFPs) were sampled and stored at 1250 Hz. Analyses were performed with custom-made Matlab programs, based on generic code that can be downloaded at http://www.battaglia.nl/computing/ and http://fmatoolbox.sourceforge.net/. Local field potentials were recorded using tungsten wires with PFA isolation (0.002” bare). Tetrodes were plated to reduce impedances to around 100 kΩ using gold solution (Neuralynx).

### Sleep scoring

Once the data had been pre-processed, an automatic analysis of vigilance states developed in the team (Bagur et al., 2018) was carried out using Matlab software. This analysis is based on the signals of the local field potentials (LFP) of the olfactory bulb (OB) and the hippocampus (HPC). Briefly, the gamma power in the OB is low during both REM and NREM sleep, allowing us to distinguish wakefulness from sleep states. NREM sleep is distinguished from REM sleep based on the theta (5-10Hz) / delta (2-4) power ratio in the HPC. Indeed, during NREM sleep there is a strong presence of slow waves (high power in the delta band), while theta oscillations are highly prominent during REM sleep.

### Histological analysis

After completion of the experiments, mice were deeply anesthetized with intraperitoneal infection of ketamine (1%) and xylazine (1‰). The animals were then perfused transcardially with saline (approximately 50 ml), followed by approximately 50 ml of PFA (4 g/100 mL). Brains were extracted and placed in PFA for postfixation for at least 24 h, transferred to sucrose for at least 48 h, and then cut into 40-μm-thick sections using a freezing microtome and mounted and stained with hard set vectashield mounting medium with DAPI (Vectorlabs)

### Data analysis and statistics

Data analysis and statistics were performed in the Matlab environment (MathWorks) and R (version 4.2.2). All values are expressed as means ± s.e.m and statistical significance on all figures uses the following convention of p-values: * p <0.05, ** p < 0.01, *** p <0.001. A normality test was performed on each dataset using the Shapiro-Wilk test. The parametric tests (paired or unpaired t-tests, one-way or two-way ANOVAs) were used if the datasets were normally distributed or if no non-parametric equivalent was available. For all optogenetic experiments, paired or unpaired t-tests were performed accordingly. Paired t-tests were also employed for the effect of the chemogenetic inhibition of the PFC-VLPO pathway under basal conditions (without SDS procedure). One-way ANOVAs or two-way repeated measures ANOVAs were performed accordingly for SDS data. ANOVAs were followed by a post-hoc Wilcoxon rank-sum or signed-rank test with a Benjamini-Hochberg correction (in this case corrected p-values are shown).

## RESULTS

### PFC sends excitatory monosynaptic projections to VLPO sleep-promoting neurons

The presence of PFC afferents in the VLPO has been previously demonstrated in rats (Chou et al., 2002). We first confirmed this anatomical pathway in mice. We injected AAV-CamK2-ChR2-mCherry into the PFC, targeting glutamatergic projection neurons expressing the CamK2 promoter (Fig. 1A). Injection sites were well centered in the infralimbic (IL) and prelimbic cortex (PrL) cortex and sometimes extended ventrally to the Taenia tecta (TT) (Fig. 1B). In the same animals, mCherry labeling was also observed locally at the level of the preoptic area where it was densely packed in the VLPO (Fig. 1C-D). This anatomical evidence demonstrates the existence of direct projections from the PFC into the VLPO in mice. In order to test the functionality of this cortical pathway, we performed *ex vivo* recordings on brain slices from mice injected with AAV-CamK2-ChR2-mCherry into the PFC (Fig. 1E-H). We identified 13 VLPO neurons that responded to optogenetic stimulation (blue light, 5ms). These neurons recorded in whole-cell, voltage-clamp mode at -60 mV displayed evoked inward currents with an average amplitude of 33.08 ± 7 pA, a duration of 25.68 ± 4.40 ms, and a latency of 2.54 ±0.22 ms (Fig. 1F). The current was always inward whatever the holding potential imposed from -55 mV to -95 mV. To test the monosynaptic nature of this current, we performed the photostimulations in the presence of voltage-gated sodium (TTX 1 µM) and voltage-gated potassium (4-AP 100 µM) channel blockers. At -60 mV, although their amplitude was decreased, these evoked currents persisted, demonstrating their monosynaptic character (n=3/3, Fig. 1F). Furthermore, application of a non-selective glutamate receptor antagonist, kynurenate (1 mM), abolished these evoked currents, confirming their glutamatergic nature (n = 3/3, Fig. 1G). To characterize the properties of these responding neurons, we recorded them in current-clamp configuration. This revealed that they mostly displayed a low threshold spike (LTS, n = 10/13, Fig. 1I) and were hyperpolarized by noradrenaline (n = 5/6, Fig. 1J). It is well established that these two properties (LTS and NA-induced inhibition) are indicative of VLPO neurons corresponding to sleep-promoting neurons recorded *in vivo* (Gallopin et al., 2000, 2005; Matsuo et al., 2003; Saint-Mleux, 2004; Liu et al., 2010). Therefore, the PFC monosynaptically excites sleep promoting neurons in the VLPO.

**Figure 1:**
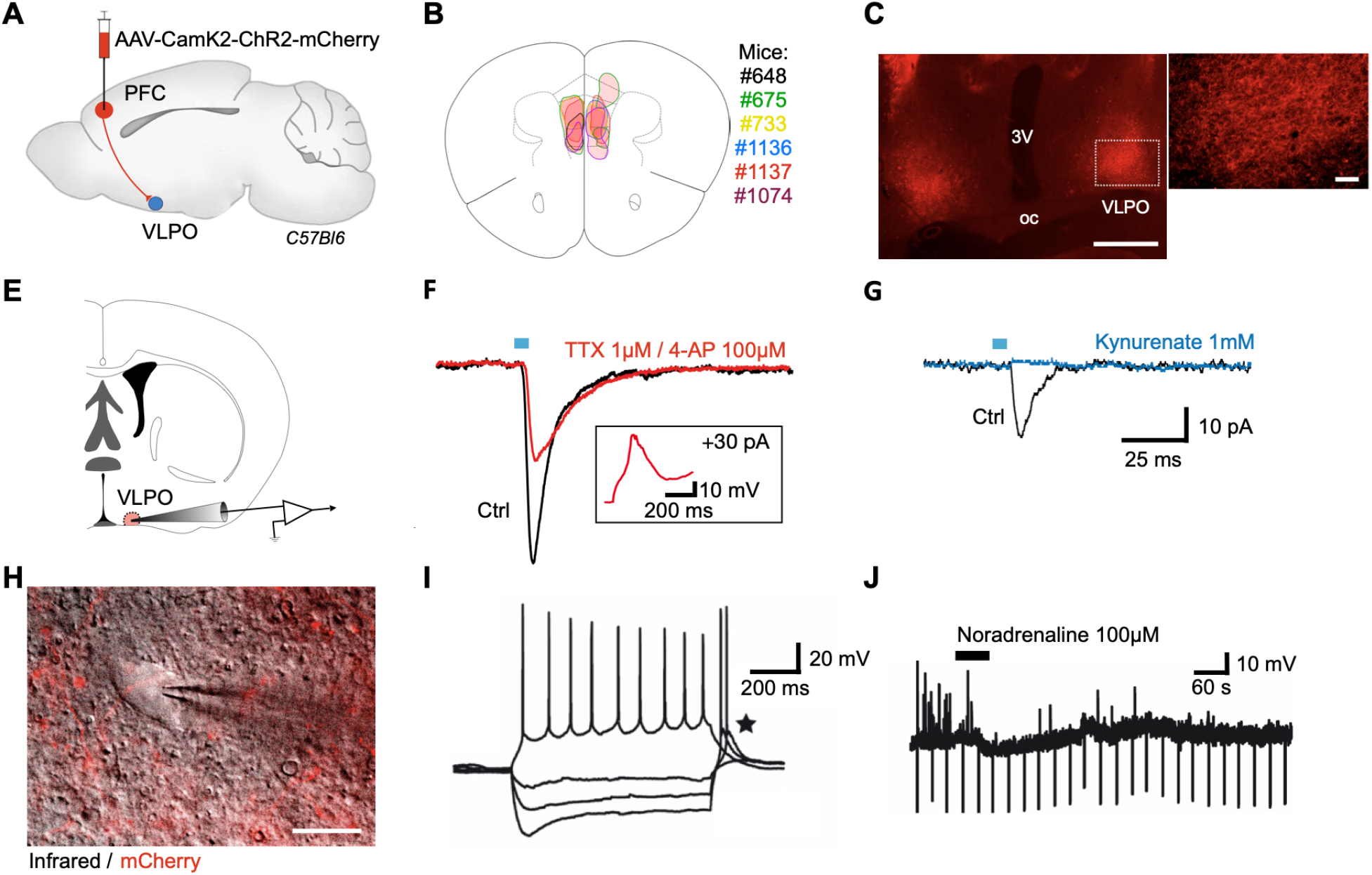
CamK2-expressing neurons in the prefrontal cortex project directly to the VLPO. **A**, Schematic representation of viral vector injections (AAV-CamK2-ChR2-mCherry) into PFC to express ChR2 in projection neurons to the VLPO. **B**, Outlines of viral injection sites in the IL/PL (n = 6 mice). **C**, Left: fluorescence microphotograph of a representative VLPO slice illustrating the dense mCherry labeling of PFC axonal terminals in VLPO (scale bar, 1 mm). Right: magnification of the area delineated by the dashed lines in C (scale bar, 100 µm). **E**, Schematic of a brain slice illustrating the *ex vivo* patch-clamp recording protocol of VLPO neurons. **F**, Average traces of currents evoked by photostimulation in control conditions (black trace) and with TTX/4AP (red trace) in a representative neuron (measured at -60 mV). The insert shows the voltage response in current-clamp mode with TTX/4AP. Note the presence of the LTS without sodium action potentials. **G**, Average traces of evoked currents in control condition (black trace) and in the presence with kynurenate (blue trace) in a representative responsive neuron. **H**, Superimposed infrared and fluorescence microphotographs of a VLPO neuron recorded in patch-clamp conditions. mCherry fluorescence reveals axonal fibers that are in contact with the neuron (scale bar, 20 µm). **I**, Discharge profile of action potentials (APs) evoked by step application of increasing currents (-100 pA, -60 pA, -20 pA, 20 pA) from the recorded neuron. Note the presence of a LTS visible at the end of the hyperpolarizing current steps (star). **J**, Induced hyperpolarization by noradrenaline of a representative cell responding to optogenetic stimulation. Each downward deflection represents the voltage response to the regular application of a hyperpolarizing current step.

### Activation of the PFC-VLPO pathway does not affect NREM sleep but suppresses REM sleep episodes and favors transition to NREM

Next, we investigated the functional role of this pathway by examining *in vivo* its impact on sleep. Wild-type mice were injected in the PFC with AAV-CamK2-ChR2-mCherry (ChR2 mice, n=6) or AAV-CamK2-mCherry as a control (mCherry mice, n=5). All animals were implanted with intracerebral electrodes to perform sleep scoring based on local field potentials (LFPs) recorded in the PFC, Olfactory Bulb (OB), hippocampus (HPC). In the VLPO bilateral optical fibers were also implanted(Fig. 2A-B). We delivered 30s light trains at 20 Hz to the VLPO iht during both NREM and REM sleep. In order to separate the effect of the activation of cortical afferents on specific sleep states, we separately analyzed the photostimulations that appeared during sustained (>5s) NREM or REM sleep episodes respectively. When given during NREM sleep, the stimulations did not induce any significant difference in sleep or wake states when comparing ChR2 mice to mCherry mice (Fig. 2C-E). This is confirmed by the analysis of average spectrograms in the PFC which show that activating cortical afferents in the VLPO did not change the power of slow oscillations (1-4 Hz) characteristic of NREM sleep (laser OFF vs laser ON; m = 7.1938 ± 1.0147, 6.8395 ± 0.9274; p = 0.2434 (ChR2); m = 9.4364 ± 1.6958, 9.4364 ± 1.6908; p = 0.3732 (mCherry), Fig. 2F-G).

**Figure 2:**
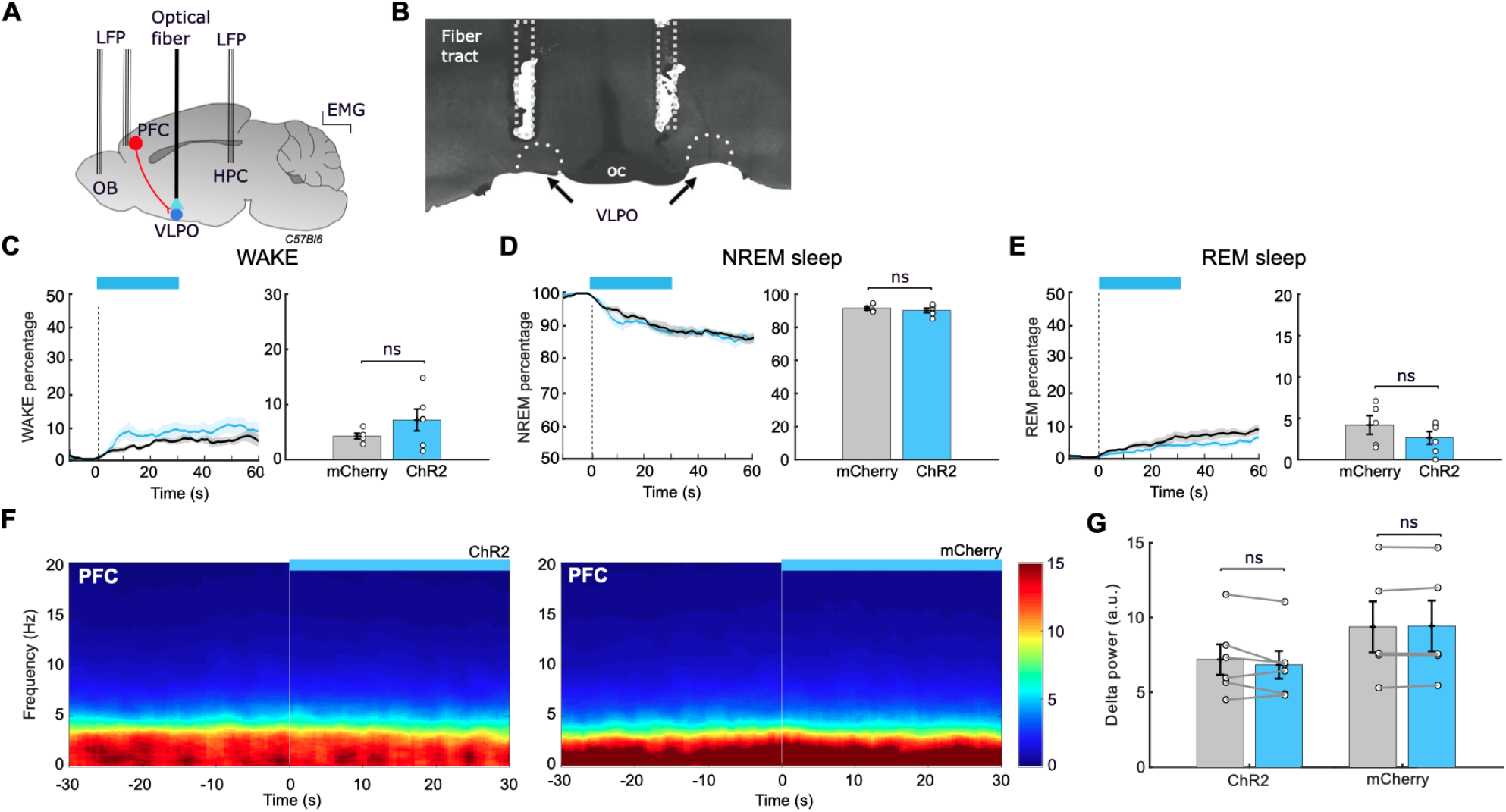
Optogenetic activation of PFC-VLPO pathway does not affect NREM sleep. **A**. Schematic representation of the electrodes implanted in the OB, HPC, PFC. Optical fibers were bilaterally implanted above the VLPO to deliver light. **B**. Representative example of a coronal slice showing the location of the optic fibers above the VLPO. **C**. Left : Percentage of wakefulness as a function of time during the photo-stimulations triggered in NREM sleep for ChR2 (blue, n=6) and mCherry (gray, n=5) mice . Right : Average wake percentage during the 30s photo-stimulation time window. **D**. Same as in C. but for NREM sleep percentage. **E**. Same as in C. but for REM sleep percentage. **F**. Representative examples of spectrograms of the PFC for a ChR2 mouse (left) and a mCherry mouse (right) during photo-stimulations triggered during NREM sleep. **G**. Quantifications of the average power in the delta band (1-4Hz) during the 30s time windows before (gray bars) and during the NREM photo-stimulations (blue bars) for ChR2 (n = 6) and mCherry (n = 5) mice. The blue rectangles represent the photo-stimulations (laser ON).

However, when stimulations were given during REM sleep episodes, we observed a decrease in the percentage of this state for the ChR2 mice compared to mCherry (Fig. 3C). The quantification of the REM percentage over the 30-seconds of stimulation confirmed the significant decrease of REM for the ChR2 compared to mCherry mice (m = 45.88 ± 6.03 (ChR2), m = 66.61 ± 3.98 (mCherry), p = 0.0229, Fig. 3C). As a mirror effect, we found that the decrease in REM sleep induced by the stimulation was compensated by a significant increase in the NREM sleep percentage in ChR2 mice compared to mCherry mice (m = 49.40 ± 5.20 (ChR2), m = 31.4926 ± 4.1771 (mCherry), p = 0.0286, Fig. 3B). The decrease in REM sleep is consistent with the analysis of spectrograms performed in the HPC, which indicated a significant decrease in theta power during the 30s stimulations as compared to the 30s prior to stimulations for the ChR2 mice (m = 3.3210 ± 0.2402 (laser OFF), 2.5695 ± 0.2840 (laser ON), p = 0.0151, Fig. 2D-E). No difference was found for the mCherry mice (m = 3.3189 ± 0.1298 (laser OFF), 3.1083 ± 0.1400 (laser ON), p = 0.0814, Fig. 2D-E). Consequently, it appears that the decrease in theta power is the result of the photoactivation of cortical afferences.

**Figure 3:**
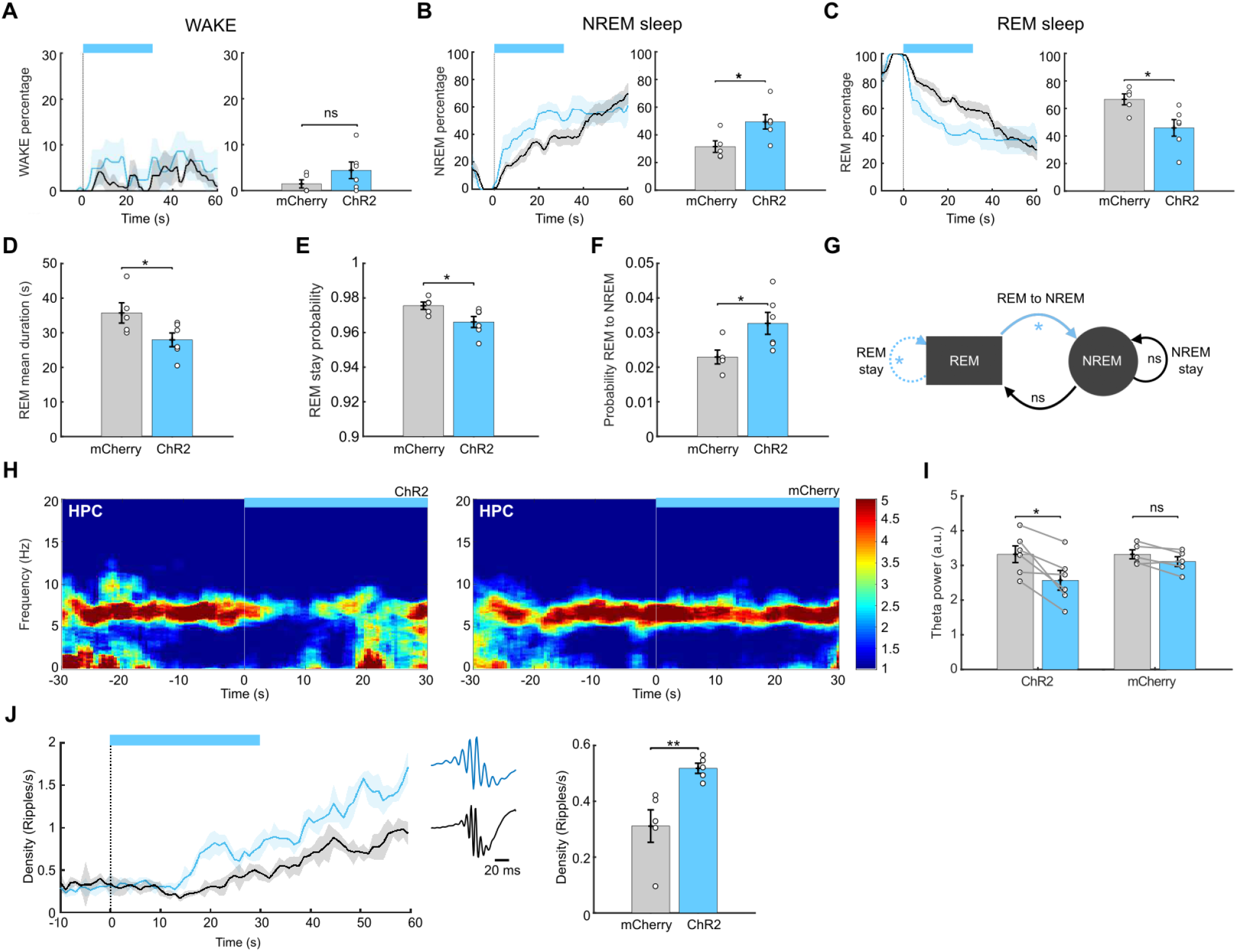
Optogenetic activation of PFC-VLPO pathway fragments REM sleep episodes and favors a transition to NREM sleep. **A**. Left : Percentage of wakefulness as a function of time during the photo-stimulations triggered in REM sleep for ChR2 (blue, n=6) and mCherry (gray, n=5) mice. Right : Average wake percentage during the 30s stimulation time window. **B**. Same as in A. but for NREM sleep percentage. **C**. Same as in A. but for REM sleep percentage. **D**. Mean duration of REM sleep bouts over the whole sleep session for ChR2 and mCherry mice. **E**. Probability to stay in REM sleep over the whole sleep session for ChR2 and mCherry mice. **F**. Probability of transition from REM to NREM sleep over the whole sleep session for ChR2 and mCherry mice. **G**. Summary scheme. The photo-activation of the PFC-VLPO pathway during REM sleep decreases the probability to stay in REM sleep and increases the probability to transition to NREM sleep, leading to REM sleep fragmentation. **H**. Representative examples of spectrograms of the HPC during photo-stimulations triggered during REM sleep for a ChR2 mouse(left) and for a mCherry mouse (right). **I**. Quantifications of the average power in the theta band (5-9Hz) during the 30s time windows before (gray bars) and during the photo-stimulations (blue bars) triggered in REM sleep for ChR2 (n = 6) and mCherry (n = 5) mice. **J**. Ripples density as a function of time during the photo-stimulations triggered in REM sleep for ChR2 mice (blue, n=5) and mCherry mice (gray, n=5) (left). Average ripples waveforms for one representative ChR2 mouse (blue) and one mCherry mouse (gray). Average quantification of the ripples density in the 30s time window during photo-stimulation triggered in REM sleep (right). Blue rectangles represent the photo-stimulations (470nm, 20Hz, 30s). Unpaired t test.

Importantly, proportions of arousal were similar in ChR2 and mCherry mice during stimulation periods, showing that the activation of cortical afferents in the VLPO does not awaken the animals (Fig. 3A). All these results suggest that optogenetic activation of the PFC-VLPO pathway during REM sleep leads to a predominant transition to NREM sleep. Accordingly, we found that the density of hippocampal sharp-wave ripples, a specific electrophysiological marker of NREM sleep was significantly enhanced in ChR2 mice compared to mCherry mice during stimulations delivered in REM sleep (m = 0.5181 ± 0.0182 (ChR2), m = 0.3112 ± 0.0582 (mCherry), p = 0.0094, Fig. 3J). These results reinforce our previous conclusions suggesting that the activation of the PFC-VLPO pathway promotes the transition from REM to NREM.

We then analyzed in more detail the effects of photo-stimulation of the PFC terminals in the VLPO on the number as well as the duration of REM episodes on the global sleep session. We noticed that there is a decrease in the REM sleep bouts mean duration in ChR2 mice compared to mCherry mice (m = 27.9533 ± 1.9826 (ChR2), m = 35.7212 ± 2.9273 (mCherry), p = 0.049, Fig. 3D). This is consistent with the fact that the stimulations delivered during REM sleep suppress the ongoing episode (on the time scale of stimuli, see above). We also quantified the transition probability between sleep states over the entire sleep session. We found that ChR2 mice had a lower probability of remaining in REM sleep compared to mCherry mice (m = 0.0996 ± 0.0032 (ChR2), m = 0.9755 ± 0.0021 (mCherry), p = 0.0434, Fig. 3E). Consequently, the activation of the PFC-VLPO pathway induces a fragmentation of REM sleep through a perturbation of REM sleep maintenance. Still in line with what we observed at the timescale of the stimulations, the decrease of REM sleep maintenance in ChR2 mice is associated with a significantly higher probability of transition from REM to NREM sleep compared to mCherry mice (m 0.0327 ± 0.0032 (ChR2), m = 0.0229 ± 0.0020 (mCherry), p = 0.0359, Fig. 3F). Altogether,our results show that the activation of the PFC-VLPO pathway decreases the duration of the REM sleep episodes and affects the maintenance of REM sleep computed over the entire sleep session. Moreover, this exit of REM sleep induced by our simulations is in favor of NREM sleep which is consistent with the NREM sleep promoting role of the VLPO.

### The pharmacogenetic inhibition of PFC-VLPO projections does not affect sleep-wake architecture under basal conditions

In light of these findings, we sought to understand the ecological context in which the PFC-VLPO pathway could decrease REM sleep. Exposure to certain uncontrollable stressors has been shown to reduce REM sleep (Sanford et al., 2005, Henderson et al., 2017). We therefore hypothesized that the PFC-VLPO pathway might be involved in this effect.

We first investigated the effect of the inhibition of the PFC neurons that project to VLPO in normal conditions. A retrovirus allowing the expression of the Cre-recombinase was injected in the VLPO of wild-type mice(rAAV-hSyn1-EGFP-Cre). A second virus (AAV-hSyn1-hM4D(Gi)-mCherry) was then injected into the IL cortex to allow the inhibitory DREADD receptor to be expressed in IL neurons that project to the VLPO (Fig. 4A). Projecting neurons could easily be identified thanks to the GFP fluorescence. We observed a multitude of labeled GFP neurons in the IL/PL cortex, confirming the results of our anterograde tracing and providing further evidence for the existence of the PFC-VLPO anatomical pathway (Fig. 4C). With confocal microscopy, we verified that the DREADD receptor (with mCherry labeling) was specifically expressed in GFP-labeled neurons. This confirms that our chemogenetic experiments will specifically target the PFC neurons that project to the VLPO (Fig. 4D-F). To confirm that the DREADD receptors were functional, the effect of CNO on the membrane potential of PFC neurons identified by mCherry fluorescence was examined *ex vivo* in PFC slices (Fig. 4G). Bath application of CNO (10 µM) systemically induced a hyperpolarization in all the mCherry expressing cells we recorded (n=4/4, Fig. 4H). The same viral strategy was then applied in mice that were implanted with intracerebral electrodes to test *in vivo* the effects of inhibiting the PFC neurons that project to the VLPO on sleep. Seven mice received intraperitoneal injections of either CNO (2,5 mg/kg) or saline. In comparison with saline-treated mice, CNO treatment did not result in any change in the proportions of wake, NREM or REM sleep (saline vs CNO; m = 21.0585 ± 1.2999, 19.8699 ± 2.2627; p = 0.6026 (wake); m = 68.2615 ± 1.4390, 70.0293 ± 2.0397; p = 0.3998 (NREM); m = 10.6800 ± 0.5635, 10.1007 ± 0.6319; p = 0.3479 (REM), Fig. 4I-K). Therefore, the inhibition of the PFC-VLPO pathway is not involved in the regulation of sleep states under baseline conditions.

**Figure 4:**
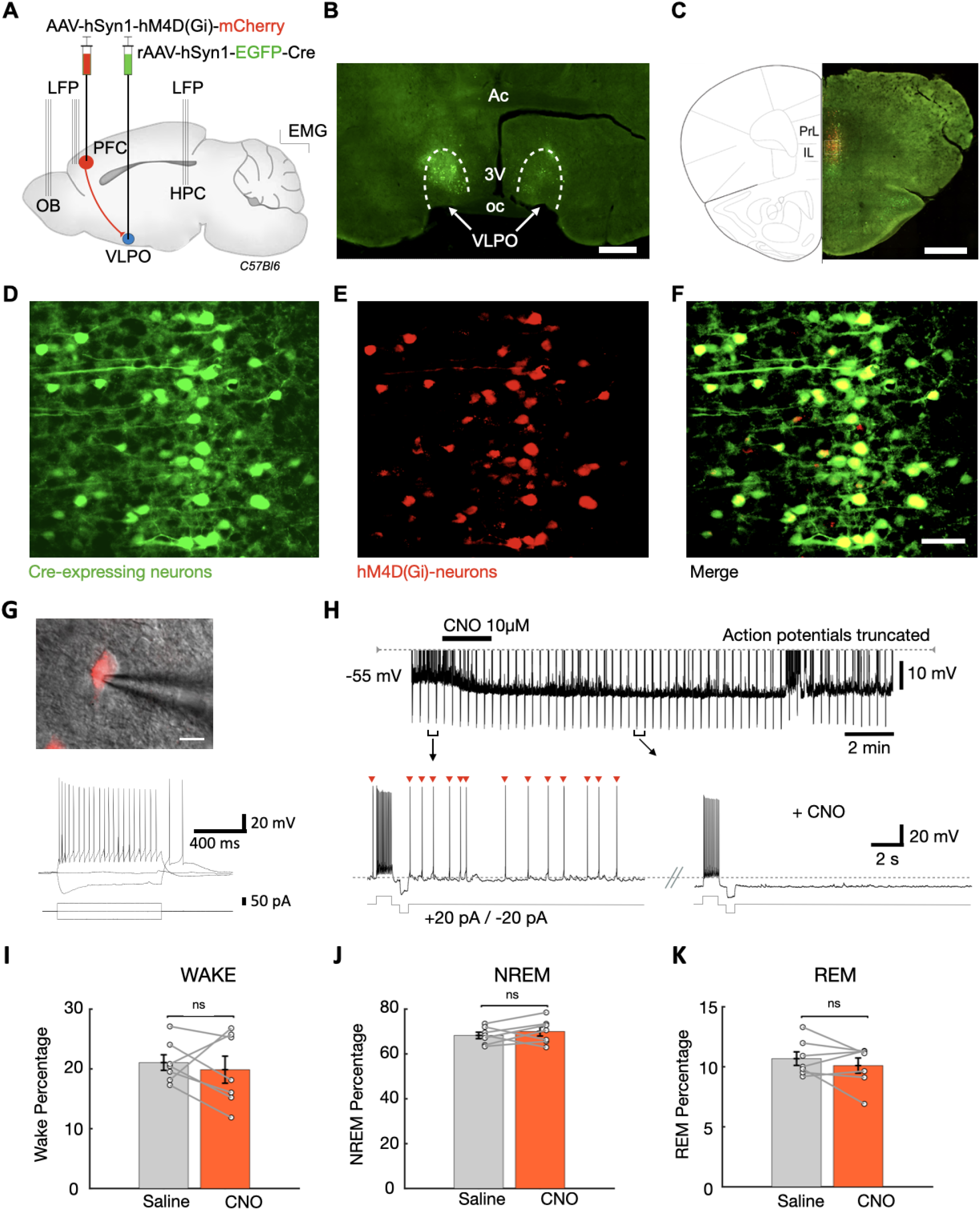
Chemogenetic Inhibition of PFC neurons projecting to the VLPO has no effect on baseline sleep-waking states. **A**, This schematic illustrates the injection of viral vectors into the VLPO (rAAV-hSyn1-EGFP-Cre) and into the PFC (AAV-hSyn1-hM4D(Gi)-mCherry) to express the hM4D receptors specifically in PFC neurons that project to the VLPO. **B**, Microphotograph of a representative slice illustrating by EGFP fluorescence the injection site in the VLPO (scale bar, 1 mm). **C**, Overlay of green and red fluorescence in a representative slice revealing the presence in the PFC of mCherry- and GFP-labeled neurons (scale bar, 1 mm). **D, E**, Microphotographs at a higher resolution showing in the PFC a dense labeling of retrogradely labeled GFP neurons expressing the Cre-recombinase **(D)** and mCherry neurons expressing the inhibitory DREADD receptors **(E). F**, The superposition of images in D. and E. reveals strong expression of hM4D receptors in retrogradely labeled neurons (scale bar, 50 µm). **G**, Superimposed infrared and fluorescence microphotographs of a PFC neuron recorded in patch-clamp conditions (scale bar, 20 µm). **H**, The upper panel illustrates a hyperpolarization of a mCherry labeled neurons following bath application of CNO (10 μM). During recording, hyperpolarizing (-20 pA) and depolarizing (-20 pA) current steps were elicited at regular intervals. The lower panels show enlargements of the voltage responses before CNO application and during the effect of the CNO. Note that the spontaneous AP at baseline (red arrowheads) disappear during the CNO effect. The AP discharge frequency is also decreased in response to the depolarizing current step during the CNO effect. **I, J K**, Illustrate the percentages of Wake, NREM and REM sleep respectively, after saline and CNO injections (i.p). (Paired t test, n=7).

**The PFC-VLPO pathway inducesREM sleep fragmentation following stressful events We next evaluated** the effect of inhibiting the PFC-VLPO pathway in a stressful situation. It has recently been demonstrated that stress induced by social defeat stress (SDS) decreases REM sleep rapidly in the hours following the event (Henderson et al., 2017). Therefore, we evaluated how this type of stress affects REM sleep. Our experimental mice were submitted to SDS by exposing them to a large, aggressive CD1 mouse. They were then treated either with CNO (2,5 mg/Kg) or saline and their sleep and wake states were monitored in their home cage for the rest of the day (Fig. 5A). Mice undergoing SDS were compared to the same mice previously measured during the same period of time without exposure to the aggressor mouse (control group).

**Figure 5:**
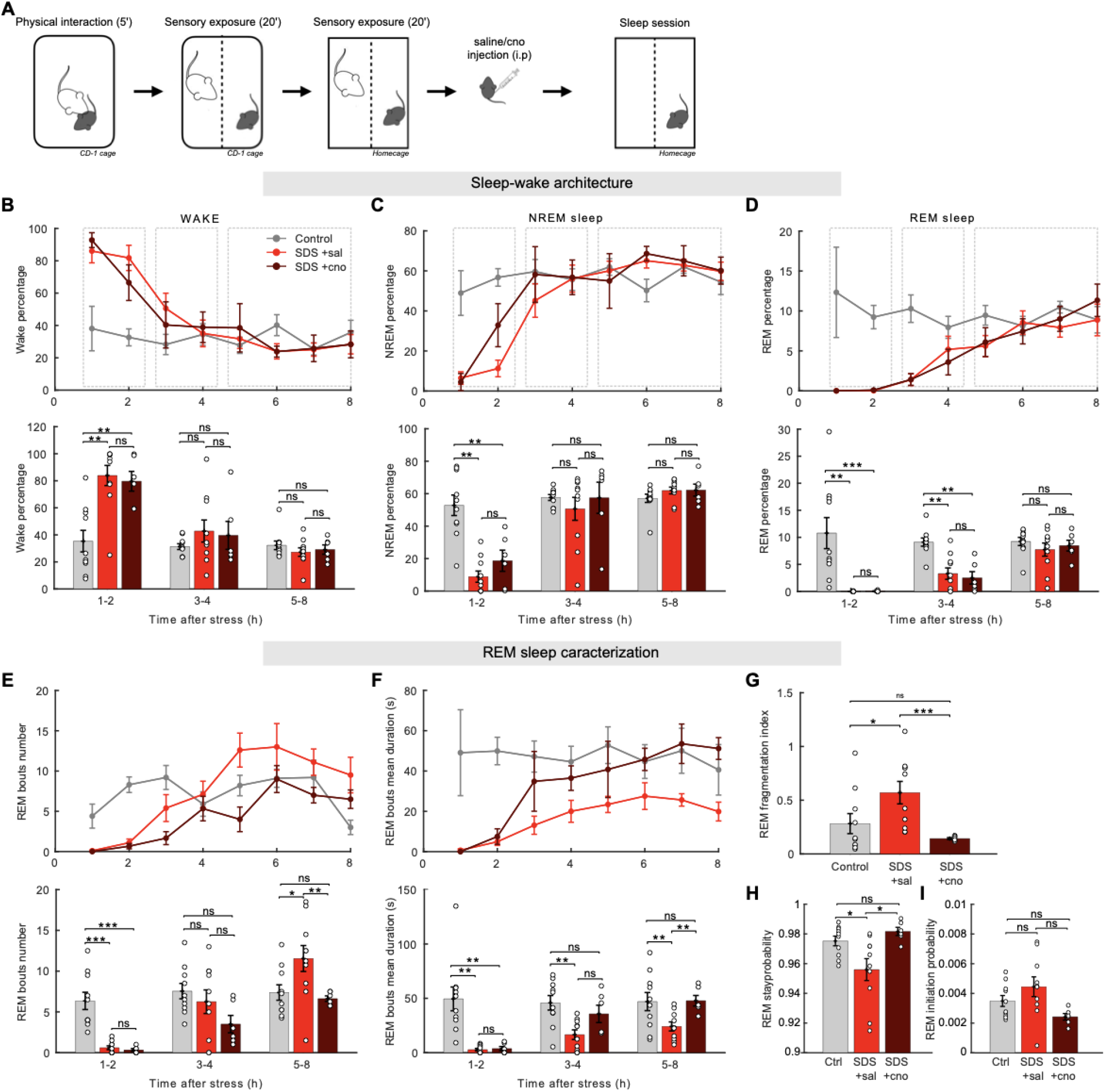
Chemogenetic inhibition of PFC-VLPO rescues the REM sleep fragmentation induced by stress. **A**, Experimental timeline for the social defeat stress (SDS) procedure. **B**,**C**,**D**, Percentage over time (top) and averaged in the 1-2h, 3-4h and 5-8h time windows (bottom) of wakefulness (B), NREM sleep (C) and REM sleep (D) in control condition (gray, n=10), after SDS + saline treatment (red, n=10) or after SDS + CNO treatment (brown, n=6). **E**, Number of REM sleep bouts over time (top) and averaged in the 1-2h, 3-4h and 5-8h time windows (bottom) for control, SDS + saline treatment and SDS + CNO treatment. **F**, Mean duration of REM sleep bouts over time (top) and averaged in the 1-2h, 3-4h and 5-8h time windows (bottom) for control, SDS + saline treatment and SDS + CNO treatment. **G**, REM sleep fragmentation index calculated as the number of REM bouts divided by their duration. **H**, Probability to remain in REM sleep in control condition or after SDS + saline treatment or SDS + CNO treatment. **I**, Probability to initiate REM sleep in control condition or after SDS + saline treatment or SDS + CNO treatment. (**B-F**, Two-way repeated measures ANOVA; **G-I**, One-way ANOVA; *p < 0.05, **p < 0.01, ***p<0.001).

We identified three main phases in the sleep-wake architecture of mice submitted to SDS (see dashed-lines rectangles, Fig. 5B-D). During the first phase (from 1 to 2 hours post SDS), SDS led to a strong insomnia characterized by a significant increase in the percentage of wake at the expense of both NREM and REM sleep (Fig. 5B-D) (control vs SDS-saline-treated, p = 0.0088). The decrease in REM sleep is thus simply due to a lack of sleep. During the second phase (3 to 4 hours post SDS), NREM of SDS mice returned to levels similar to those observed in control mice (Fig. 5C). However, the amount of REM sleep remained lower in SDS mice, showing that, in this same time period SDS induced a selective suppression of REM sleep (p = 0.0029, Fig. 5D).

During the third phase (5 to 8 hours post SDS) the percentage of both NREM and REM sleep were no longer different from control mice (Fig. 5C-D), suggesting that SDS mice had recovered normal sleep. However, when we examined REM sleep states in more detail, SDS mice showed highly fragmented REM sleep. This was characterized by a significant increase in the number of REM sleep bouts (p = 0.0464, Fig. 5E) and a decrease in the duration of REM sleep bouts (p = 0.0059, Fig. 5F). We computed a fragmentation index (FI) of REM sleep by dividing the number of REM bouts by their duration. The FI was significantly higher for SDS mice 5 to 8 hours after SDS when compared to control mice (p = 0.0468, Fig. 5G).

As in our previous experiment, we quantified the probability of initiating and remaining in REM sleep (Fig3E-F). In line with the REM sleep fragmentation, we showed that the probability of staying in REM sleep decreased in SDS mice compared to control (p = 0.0107) while the probability of initiating REM sleep did not change (Fig. 5H-I). These results are reminiscent of the ones observed after the activation of the PFC-VLPO pathway (Fig3E-F). We therefore looked at the effect of the PFC-VLPO pathway inhibition on sleep after SDS. At first sight, when examining the global sleep-wake architecture in terms of sleep states percentage, we did not observe any significant differences between the SDS-saline-treated group and the SDS-CNO-treated group (Fig. 5B-D). However, when looking at the transition between sleep states, we observed that the inhibition of the PFC-VLPO pathway after SDS totally reversed the REM sleep fragmentation induced by SDS. As a result, in the 5 to 8 hours period after SDS, the mean number of REM sleep bouts (Fig. 5E) as well as their mean duration (Fig. 5F) were no longer different from control (p = 0.9371, p = 0.8749, respectively). Similarly, the FI of SDS-cno-treated mice was also no longer different from control mice (p = 1), meaning that inhibition of the PFC-VLPO pathway prevented the REM sleep fragmentation induced by SDS (Fig. 5G). Finally, we found that the inhibition of the PFC neurons that project to the VLPO restored the probability of staying in REM sleep after SDS (p = 0.5742, Fig. 5H). Altogether, our results show that the PFC-VLPO pathway plays an important role in modulating maintenance of REM sleep but not its initiation, a result consistent with our experiments using the optogenetic activation of the PFC-VLPO pathway. Moreover, our results suggest that this pathway could be recruited under stressful situations in order to disrupt the REM sleep maintenance.

## DISCUSSION

A major challenge in sleep research is understanding the mechanisms that regulate the alternation between different sleep states and wakefulness. In the present study, we characterize a new neuronal pathway involving a high level structure, the prefrontal cortex, that is involved in regulating stress-induced changes in sleep patterns. We first demonstrated in mice that the prefrontal cortex sends excitatory monosynaptic projections to VLPO sleep-promoting neurons. The *in vivo* photoactivation of cortical axonal fibers in the VLPO suppressed REM sleep episodes in favor of NREM sleep. We also provide evidence that this pathway is necessary to induce the REM sleep fragmentation that we identified as part of the response to social stress.

### VLPO sleep-promoting neurons receive excitatory monosynaptic projections from the PFC

Based on both anterograde and retrograde tracing performed with viral vectors, we found that VLPO receives direct projections from the Infralimbic (IL) / Prelimbic (PrL) part of the PFC, confirming anatomical evidence previously described in rat (Chou et al., 2002). *Ex vivo* photoactivation of these cortical afferences revealed inward glutamatergic currents in VLPO neurons. The amplitude and duration of these currents are highly consistent with their glutamatergic nature (Schöne and Burdakov, 2012; Chee et al., 2015). The short latency of their appearance after photostimulation and their persistence under conditions of synaptic uncoupling (TTX-4AP) (Petreanu et al., 2007) confirm the monosynaptic nature of this connexion (Piñol et al., 2012; Rajasethupathy et al., 2015). We also demonstrated that neurons responding with this excitatory current displayed a LTS and were inhibited by noradrenaline, two properties associated with the NREM-active neurons recorded *in vivo* (Gallopin *et al*., *2000, 2005)*.

### Photo-activation of the PFC-VLPO pathway suppresses REM sleep episodes and promotes the transition to NREM sleep

Here we demonstrated that stimulations of cortical afferents in the VLPO only elicited an effect when occurring during REM sleep episodes. This effect was associated with a decrease in REM sleep compensated by an increase in NREM sleep, which is consistent with VLPO’s role in promoting NREM sleep (Saper et al., 2001). Further support for these findings can be found in the demonstration that photostimulations also increased the density of Sharp wave–ripple (SPW-R) that appeared during the induced NREM episodes. Previous research has demonstrated that a lesion in the ventromedial part of the PFC, including the IL, increased the quantity and fragmentation of REM sleep in rats (Chang et al., 2014). In light of these findings, it appears that there is a strong link between the PFC and REM sleep. Combined with our findings, we strongly believe that the modulation of REM sleep by the PFC involves at least in part an influence on the VLPO neuronal activity. Importantly, although PFC appears to play a role in the regulation of REM sleep, this pathway is unlikely to be functional in the basal condition. Indeed when we inhibited the PFC neurons that project to the VLPO, baseline sleep was not changed. Therefore the top-down cortical projections are likely recruited to the sleep network only under certain conditions.

### REM sleep is reduced by the PFC-VLPO pathway during stressful circumstances

We then raised the question of the role of this PFC-VLPO pathway and under what circumstances it can decrease REM sleep. In a stressful condition related to the threat of predation, rodents are expected to be more vigilant to increase their probability of survival. Among the sleep states, REM sleep has the highest awakening threshold and is therefore the riskiest state for many preys (Dillon and Webb, 1965; Van Twyver and Garrett, 1972; Neckelmann and Ursin, 1993). We can therefore hypothesize that the activation of the PFC-VLPO pathway would have a protective role in a stressful situation by decreasing REM sleep and privileging NREM sleep (Lima et al., 2005). We thus investigated whether inhibiting PFC neurons that project to the VLPO could prevent the decrease in REM sleep that occurs during stress. Here we tested a stress induced by social defeat well known to modulate the amount of REM sleep. It is important to note that the effects of this type of stress on REM sleep vary according to the time of day it is initiated, the duration of the exposure to the aggressive mouse, and the chronicity or acuteness of the stress experience (Henderson et al., 2017; Wells et al., 2017; Yu et al., 2022). We confirmed the result of Henderson et al that REM sleep decreases after the stress protocol. Moreover, we found that the duration of REM episodes was decreased while the number of episodes was increased. These results differ somewhat from those obtained by Henderson et al, who reported that the REM sleep reduction was mediated by a decrease in the number of REM sleep episodes and a decrease in their mean duration. Our difference could be attributed to the fact that in our conditions, the stress is prolonged by a second sensory exposure with the aggressor in the tested mouse’s cage in order to simulate a predatory environment within its enclosure. The decrease in the duration of REM sleep episodes that we observed indicates that the stress induced in our conditions would mainly affect the maintenance of REM sleep.

In comparison with control mice, we observed similar tendencies in the total number of vigilance states of mice exposed to SDS and treated either with Saline or CNO. We distinguished three main phases. During the first phase (from 1 to 2 hours after SDS), mice that had experienced SDS suffered from a strong insomnia. During the second phase (from 3 to 4 hours after SDS), mice began to sleep normally, and the amount of NREM sleep returned to normal levels. However, the percentage of REM sleep remained very low. During this second phase, there is a selective suppression of REM sleep, which is consistent with previous studies showing significant reductions in REM sleep after SDS during the first three hours (Henderson et al., 2017). The amount of sleep (both NREM and REM sleep) returned to control levels during the third phase, and the mice appeared to sleep normally thereafter. However, when we examined sleep periods more closely, we noticed that mice that experienced SDS had highly fragmented REM sleep. This was visible only when investigated simultaneously, the effects of the number and the duration of REM sleep episodes.

Our analyses indicated that the chemogenetic inhibition of PFC neurons that project to the VLPO following the stress event prevented the decrease in REM episodes, thus supporting our hypothesis that the PFC-VLPO pathway would serve as a protective mechanism in preventing the animal from staying too long in a REM period in a risky environment (Lima et al., 2005). According to our probability analysis, the animal would tend to exit REM episodes more frequently. Fragmenting REM episodes in a stressful environment instead of simply suppressing REM sleep could be an advantageous strategy since it balances environmental constraints with the need for sleep. On the one hand, it avoids long consecutive periods of unresponsiveness during REM, thus allowing the animal to monitor the environment at regular intervals. On the other hand, it preserves the total amount of REM thus conserving the beneficial functions of this sleep phase. Notably, when total sleep duration was considered, inhibition of PFC projection neurons was not able to prevent stress-induced REM loss observed early after the stressful experience. Therefore, more than just the activation of the PFC-VLPO pathway could be responsible for the inhibition of REM after social defeat stress (Henderson et al., 2017). It is also possible that the observed increase in the number of REM episodes may be responsible for the lack of REM restoration. As a compensatory mechanism, this opposite effect may offset the increase in REM sleep duration.

### Hypotheses regarding the mechanisms underlying the alteration of REM sleep during stressful conditions

Sleep-promoting neurons of the VLPO are well known for inhibiting wake-active structures during non-REM sleep (Saper et al., 2001). Considering our results, we can speculate that these neurons would also inhibit REM active structures in order to interrupt or prevent REM episodes. Indeed, it has been shown that the VLPO sleep-active neurons send projections to the lateral hypothalamic region containing neurons expressing the melanin concentrating hormone (MCH) peptide (Cvetkovic et al., 2003). These neurons have been described to be active during REM sleep and to project to vlPAG (Luppi et al., 2012). They would allow the initiation of REM sleep through their inhibitory projections onto vlPAG REM-off neurons. In addition, GABA has been shown to hyperpolarize MCH neurons ex vivo (Gao et al., 2003). It is therefore tempting to propose that activation of GABAergic neurons in the VLPO, by photo-stimulations of cortical afferents, could inhibit MCH neurons leading to a lifting of inhibition on REM-off neurons in the vlPAG. The latter could again inhibit the REM-on neurons of the sublaterodorsal nucleus (SLD) and thus terminate the ongoing REM episode (Sakai et al., 2001; Lu et al., 2006; Hsieh et al., 2011; Weber et al., 2018; Luppi and Fort, 2019). In order to confirm the contribution of this inhibitory pathway, further experiments are required, but these are beyond the scope of this present study.

### Conclusion

In the present study, we found that the prefrontal cortex is capable of regulating sleep state through the activation of VLPO sleep-promoting neurons. We propose that this neuronal pathway is specifically recruited in stress conditions during which it fragmentsREM sleep. This is consistent with the fact that REM sleep has the highest wake-up threshold in mice, making it the most vulnerable sleep state for many prey species.

## Acknowledgments

This work was supported by the Centre National de la Recherche Scientifique (CNRS) and ESPCI-Paris.

The authors declare that no competing interests exist.

